# Molecular probes of spike ectodomain and its subdomains for SARS-CoV-2 variants, Alpha through Omicron

**DOI:** 10.1101/2021.12.29.474491

**Authors:** I-Ting Teng, Alexandra F. Nazzari, Misook Choe, Tracy Liu, Matheus Oliveira de Souza, Yuliya Petrova, Yaroslav Tsybovsky, Shuishu Wang, Baoshan Zhang, Mykhaylo Artamonov, Bharat Madan, Aric Huang, Sheila N. Lopez Acevedo, Xiaoli Pan, Tracy J. Ruckwardt, Brandon J. DeKosky, John R. Mascola, John Misasi, Nancy J. Sullivan, Tongqing Zhou, Peter D. Kwong

## Abstract

Since the outbreak of the COVID-19 pandemic, widespread infections have allowed SARS-CoV-2 to evolve in human, leading to the emergence of multiple circulating variants. Some of these variants show increased resistance to vaccines, convalescent plasma, or monoclonal antibodies. In particular, mutations in the SARS-CoV-2 spike have drawn attention. To facilitate the isolation of neutralizing antibodies and the monitoring the vaccine effectiveness against these variants, we designed and produced biotin-labeled molecular probes of variant SARS-CoV-2 spikes and their subdomains, using a structure-based construct design that incorporated an N-terminal purification tag, a specific amino acid sequence for protease cleavage, the variant spike-based region of interest, and a C-terminal sequence targeted by biotin ligase. These probes could be produced by a single step using in-process biotinylation and purification. We characterized the physical properties and antigenicity of these probes, comprising the N-terminal domain (NTD), the receptor-binding domain (RBD), the RBD and subdomain 1 (RBD-SD1), and the prefusion-stabilized spike ectodomain (S2P) with sequences from SARS-CoV-2 variants of concern or of interest, including variants Alpha, Beta, Gamma, Epsilon, Iota, Kappa, Delta, Lambda, Mu, and Omicron. We functionally validated probes by using yeast expressing a panel of nine SARS-CoV-2 spike-binding antibodies and confirmed sorting capabilities of variant probes using yeast displaying libraries of plasma antibodies from COVID-19 convalescent donors. We deposited these constructs to Addgene to enable their dissemination. Overall, this study describes a matrix of SARS-CoV-2 variant molecular probes that allow for assessment of immune responses, identification of serum antibody specificity, and isolation and characterization of neutralizing antibodies.

## Introduction

The outbreak of the COVID-19 pandemic caused by the severe acute respiratory syndrome coronavirus-2 (SARS-CoV-2) has led to more than 260 million confirmed cases and 5.2 million deaths worldwide since its onset in December 2019 (https://covid19.who.int). With joint efforts by public health authorities and scientists, COVID-19 vaccines have been developed at an unprecedented speed, with several vaccines now licensed or granted emergency use authorizations. Despite the rollout of effective vaccines, widespread infections have led to the emergence of numerous variants, including multiple variants of concern (VOC) that displaced the original strain or early circulating strains around the world. These variants harbor mutations in the spike protein, some of which are associated with increased transmissibility and/or immune escape. For example, the D614G mutation contributes a fitness advantage to SARS-CoV-2 and is hence associated with enhanced infectivity [1–3], whereas the L452R, S477N, and E484K mutations may lead to reduced protection from re-infection or increased resistance to vaccine-elicited antibodies [4, 5]. Though COVID-19 vaccines approved and authorized for use thus far are still effective against these variants in preventing severe disease, there have been reports that the B.1.351 variant (Beta), the B.1.617.2 variant (Delta), and the B.1.1.529 variant (Omicron) are more resistant to neutralization by convalescent plasma, monoclonal antibody treatments, and/or vaccine-elicited antibodies than the original WA-1 strain or the previously prevalent B.1.1.7 variant (Alpha) [6–13].

Biotin-labeled molecular probes are pivotal to antibody discovery and immune evaluation. To respond to global efforts combating the SARS-CoV-2 virus, we previously designed and produced constructs incorporating an N-terminal purification tag, a site-specific protease-cleavage site, the antigen of interest, and a C-terminal sequence targeted by biotin ligase that allow for tag-based purification and in-process biotinylation. Through this strategy, we have produced wildtype SARS-CoV-2 spike ectodomain and subdomain probes allowing for the identification of potent neutralizing antibodies and nanobodies that target the receptor binding domain (RBD) or the N-terminal domain (NTD) of SARS-CoV-2 [14–19], the characterization of antibody binding affinities and specificities, and the quantification of immune responses against spike and its subdomains in nonhuman primates to inform vaccine development [20, 21] and to find correlates of elicited responses with neutralization [22]. Moreover, with the rise of SARS-CoV-2 VOCs, we found that probes incorporating mutations defined by the variants could be helpful in the characterization of the impact of VOC mutations on vaccine effectiveness [23–25].

In this study, we report production and characterization of molecular probes of spike and subdomains for SARS-CoV-2 variants. By using the strategy for wildtype probes described previously [19], we created molecular probes comprising the N-terminal domain (NTD), the receptor-binding domain (RBD), the RBD with spike subdomain 1 (RBD-SD1), and the prefusion-stabilized spike ectodomain (S2P) of diverse variants, including Alpha, Beta, Gamma, Epsilon, Iota, Kappa, Delta, Lambda, Mu, and Omicron. We characterized the physical properties of the variant probes, their antibody-binding specificity, and their capacity to sort yeast cells expressing Fab region of SARS-CoV or SARS-CoV-2 spike-binding antibodies. Finally, we examined binding of human antibodies to RBD and NTD domains from five high-profile variants through sorting B-cell libraries from two convalescent donors. Overall, this study describes how structure-based design enables advantageous production of SARS-CoV-2 variant probes that provide molecular insight into immunogenicity of SARS-CoV-2 variants when isolating neutralizing antibodies, investigating antibody specificity, and monitoring longitudinal vaccine effectiveness against emerging variants.

## Results

### Probe construct design for tag-based purification with in-process biotinylation

To make the molecular probes for the SARS-CoV-2 variants readily available, we incorporated variant mutations (as listed in Figs 1 and 2) into the probe construct we reported previously [19]. The construct design comprised three segments. The first segment was an N-terminal purification tag denoted as ‘scFc3C’, which contained a modified ‘knob-in-hole’ single chain constant portion of an antibody (scFc) followed by a cleavage site (LEVLFQGP) for the highly specific human rhinovirus 3C (HRV3C) protease [26]. This ‘knob-in-hole’ feature avoids formation of intermolecular Fc dimer and is defined by a protruding tyrosine in the N-terminal-half of the Fc-interface (T366Y), a recessed threonine in the C-terminal-half of the Fc-interface (Y407T), in conjunction with an insert of a 20-residue-linker (GGGGS)_4_ (Kabat antibody residue numbering [27]). The second segment was the sequence of interest from SARS-CoV-2 variants. We designed molecular probes of the entire spike ectodomain (residues 14 to 1208) with prefusion-stabilizing mutations (S2P), N-terminal domain (NTD), receptor-binding domain (RBD), and RBD plus subdomain 1 (RBD-SD1). The third segment, denoted as ‘10lnQQ-AVI’, comprised a 10-residue linker followed by a biotin ligase-specific sequence (GLNDIFEAQKIEWHE) at the C terminus.

**Fig 1.**
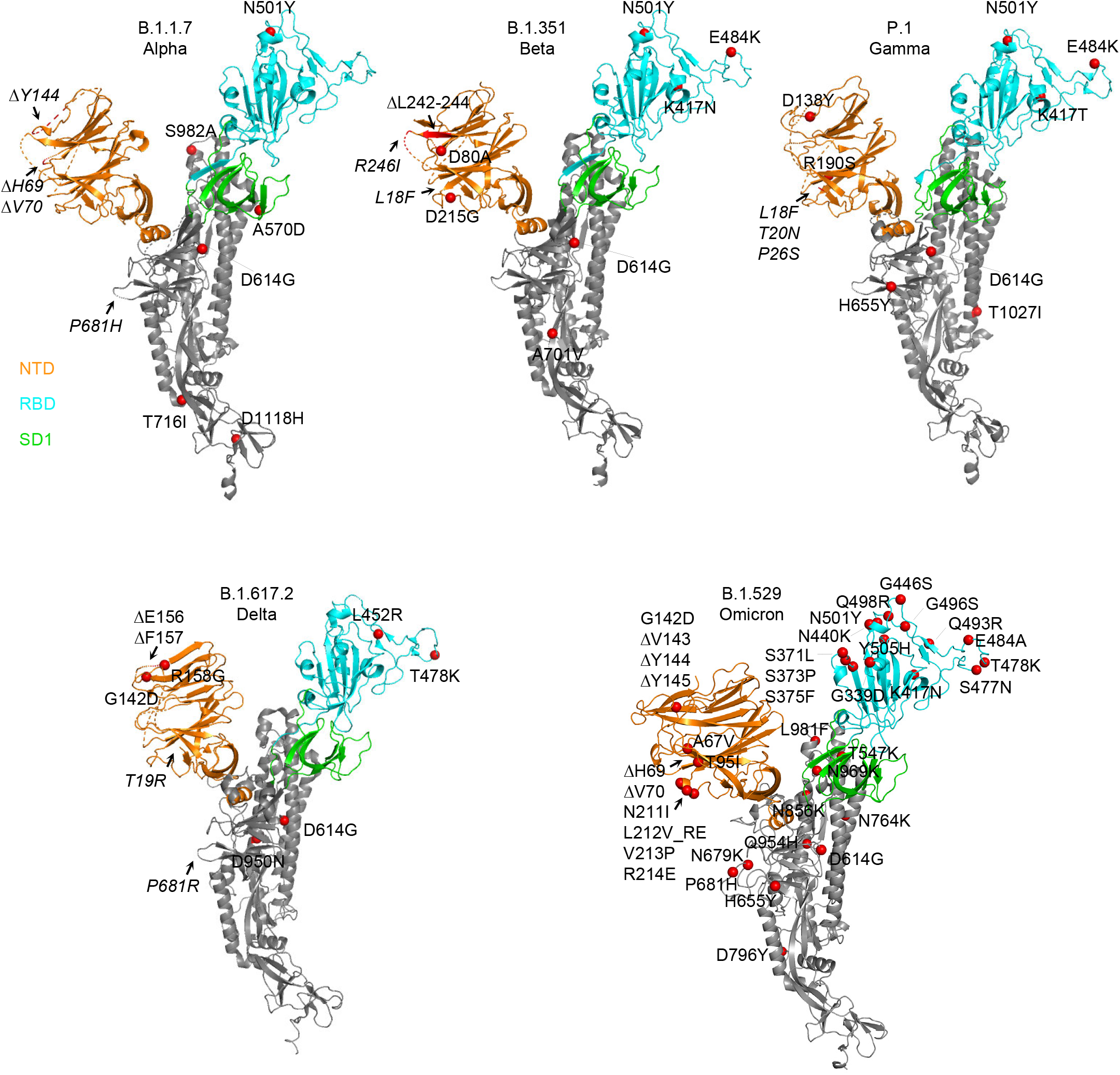
Structural modeling reveals a persistent D614G mutation, shared RBD mutations, and distinctive NTD mutations in five variants of concern. Structural models of a SARS-CoV-2 spike protomer from B.1.1.7, B.1.351, P.1, B.1.617.2, and B.1.1.529 variants. NTD, RBD, and SD1 domains are shown in orange, cyan, and green, respectively. Mutations are highlighted in the structural diagrams with a red sphere at Cα. Mutations at positions that are not resolved in the cryo-EM structure are labeled in italics with arrows pointing to the disordered regions (shown with dashed lines).

**Fig 2.**
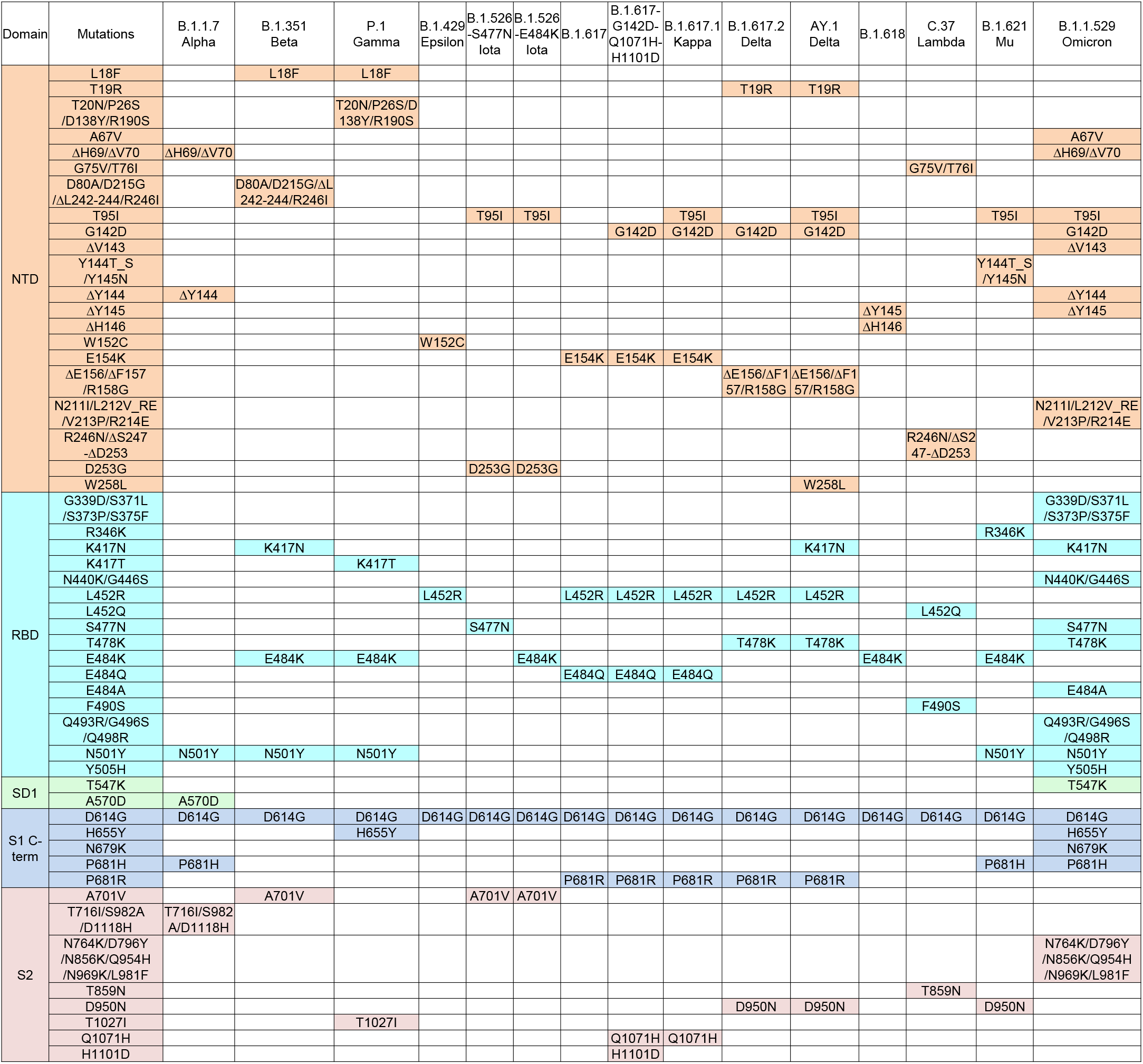
Overview of spike ectodomain mutations in variants of SARS-CoV-2. Mutations in the ectodomain of the spike protein from variants B.1.1.7, B.1.351, P.1, B.1.429, B.1.526 (2 sequences), B.1.617 (2 sequences), B.1.617.1, B.1.617.2, AY.1, B.1.618, C.37, B.1.621, and B.1.1.529 are labeled and color-coded by domain. *Note: L5F is not included in the B.1.526 sequences because the constructs in this study started from Q14.

The scFc tag allows transiently expressed proteins to be efficiently captured by protein A resins. The captured target protein can then be cleaved from the resin and biotinylated concurrently to enable streamlined purification and biotinylation on a single protein A column. This method avoids elution with acidic buffer that might alter the conformation and minimizes the need for buffer exchange and size exclusion chromatography. The plasmids for the probe constructs from this study have been deposited to Addgene (www.addgene.org) with accession numbers listed in S1 Table.

### Molecular probes of di-proline-stabilized SARS-CoV-2 variant spikes

To obtain probes of the trimeric spike ectodomain for SARS-CoV-2 variants, we incorporated the prefusion-stabilizing di-proline mutations previously reported into SARS-CoV-2 spike protein residues 14 to 1208 [28] along with a T4-phage fibritin trimerization domain (foldon) at the C-terminus [29, 30]. Briefly, the RRAR furin cleavage site at S1/S2 was replaced with GSAS, and two residues between HR1 and CH domains were substituted with prolines (K986P and V987P). The spike trimer probe constructs (hereafter referred to as ‘S2P’) were expressed by transient transfection in FreeStyle 293-F cells. In this study, we report production of S2P probes for variants B.1.1.7, B.1.351, P.1, B.1.429, B.1.526 (2 sequences; denoted as ‘B.1.526-S477N’ and ‘B.1.526-E484K’), B.1.617 (2 sequences; denoted as ‘B.1.617’ and ‘B.1.617-G142D-Q1071H-H1101D’), B.1.617.1, B.1.617.2, AY.1, B.1.618, C.37, B.1.621, and B.1.1.529. Soluble S2P trimers were captured by protein A resin through N-terminal scFc tag, followed by HRV3C-protease cleavage and BirA-catalyzed biotinylation, and finally passed through size-exclusion chromatography in PBS to remove reaction mixture and undesirable heterogeneous proteins. The final yield of the purified S2P probes ranged from ∼0.5 mg to ∼2 mg per liter of cell culture.

The purified variant S2P probes, along with WA-1 S2P and D614G S2P probes, all showed primarily a single peak by size-exclusion chromatography (Fig 3A). Size homogeneity was also confirmed by SDS-PAGE, exhibiting one major band (Fig 3B). Bio-layer interferometry (BLI) with streptavidin biosensors confirmed the biotinylation of the variant S2P proteins and ACE2 receptor binding, indicating that the variant S2P probes were well-biotinylated and recognized by ACE2 receptor (Fig 3C).

**Fig 3.**
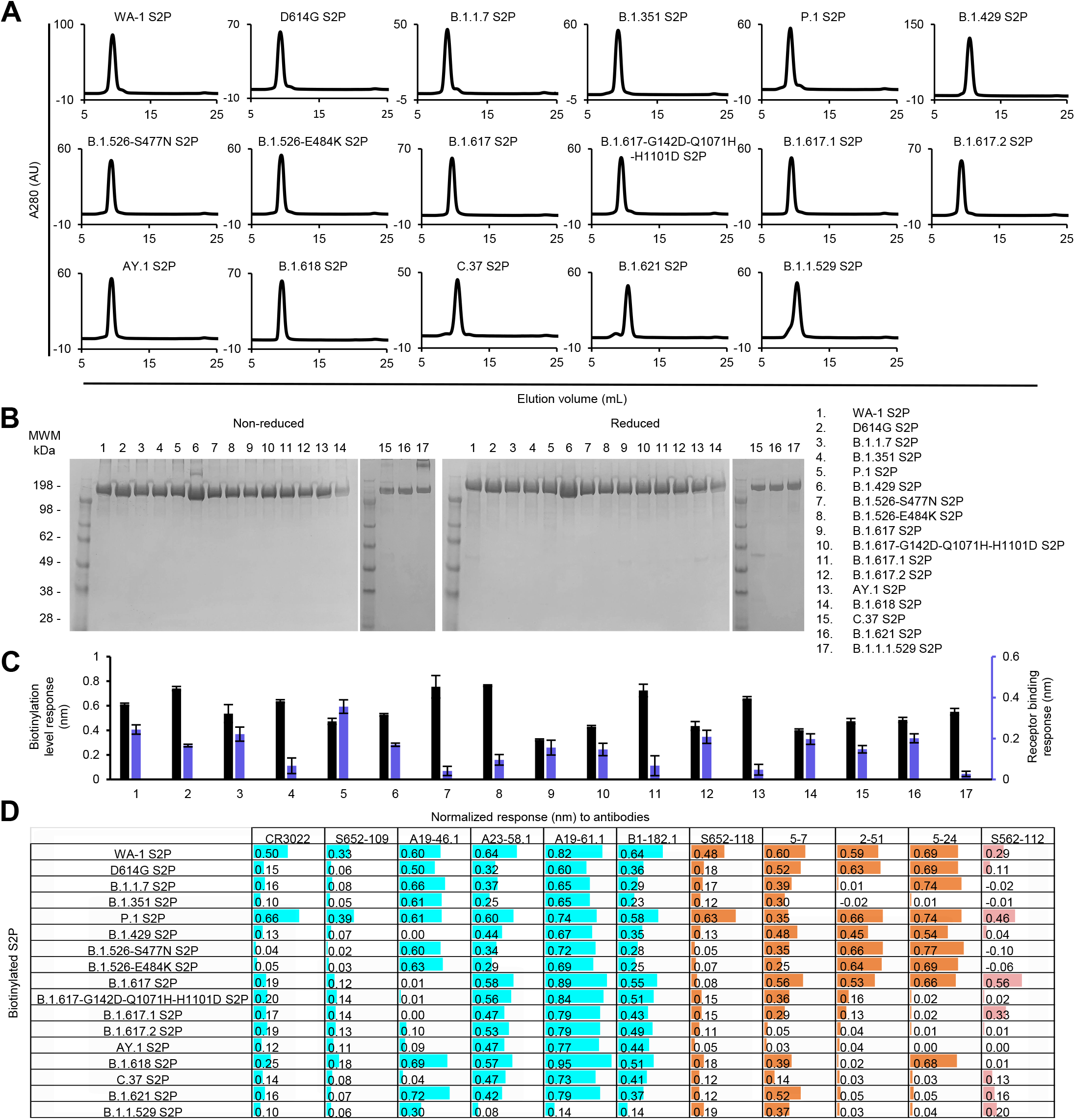
Characterization of purified biotinylated SARS-CoV-2 variant S2P probes confirms their homogeneity and antibody-binding specificity. (A) Size exclusion chromatography of the purified biotinylated SARS-CoV-2 S2P probes, including original WA-1 S2P, single mutated D614G S2P, and 15 other variant S2P, in PBS buffer. (B) SDS-PAGE of SARS-CoV-2 S2P variant probes with and without reducing agent. Molecular weight marker (MWM), the WA-1 S2P probe, and the D614G S2P probe, were run alongside the variant S2P probes. (C) Biotinylation and receptor recognition of the SARS-CoV-2 variant S2P probes. The level of biotinylation was evaluated by capture of the S2P probes at 10 μg/ml onto the streptavidin biosensors using Octet. Receptor binding was assessed with 500 nM of dimeric ACE2. Error bars represent standard deviation of triplicate measurements. (D) Antigenicity assessment of the SARS-CoV-2 variant S2P probes. Responses to RBD-directed, NTD-directed, and S2 subunit-directed antibodies were shown in cyan, orange, and rose, respectively. Bar scale between 0 and 1. Negative values not depicted.

We then characterized by BLI the binding of variant S2P probes to a panel of antibodies, including four antibodies (CR3022, S652-109, S652-118, and S562-112 [31, 32]) from SARS-CoV convalescent donors and seven antibodies (A19-46.1, A23-58.1, A19-61.1, B1-182.1, 5-7, 2-51, and 5-24 [18, 33, 34]) from SARS-CoV-2 convalescent donors (Fig 3D). The four antibodies identified from SARS-CoV convalescent donors bound to the WA-1 SARS-CoV-2 S2P. However, they showed reduced responses to many of the variant spikes. For example, the RBD-binding neutralizing antibody CR3022 and the RBD-binding non-neutralizing antibody S652-109 showed lower responses to all the S2P variants except for P.1 S2P than to WA-1 S2P. A similar pattern was observed for the NTD-specific neutralizing antibody S652-118. Unexpectedly, S652-112, which recognizes an epitope in S2, showed distinct BLI responses to three variants that have no mutations in their S2 subunit (B.1.429, B.1.617, and B.1.618); it bound B.1.617 S2P but not the other two. Among the four RBD-targeting antibodies isolated from SARS-CoV-2 convalescent patients, A23-58.1, A19-61.1, and B1-182.1 bound to all the investigated variant S2P probes except for low responses to B.1.1.529, whereas A19-46.1 recognized only S2P probes without L452 mutation. Low to no binding was observed for antibody A19-46.1 and S2P probes containing L452R or L452Q, including variants B.1.419, B.1.617, B.1.617-G142D-Q1071H-H1101D, B.1.617.1, B.1.617.2, AY.1, and C.37. Among the three NTD-directed neutralizing antibodies isolated from SARS-CoV-2 convalescent donors, 5-7 could recognize most of the variants well except two of the Delta variants, B.1.617.2 and AY.1, and the Lambda variant C.37. Antibody 2-51, which targets an undetermined region on the spike trimer but can be clustered together with other NTD-directed antibodies [34], showed impaired responses to many of the variant S2P probes, including those of B.1.1.7, B.1.351, B.1.617-G142D-Q1071H-H1101D, B.1.617.1, B.1.617.2, AY.1, B.1.618, C.37, B.1.621, and B.1.1.529. Antibody 5-24 could not recognize B.1.351, C.37, B.1.621 S2P and any variants with G142D mutation, such as B.1.617-G142D-Q1071H-H1101D, B.1.617.1, B.1.617.2, AY.1, and B.1.1.529. Notably, though these three NTD-directed antibodies showed distinct binding profiles, they all lost binding to B.1.617.2 and AY.1 S2P.

Additionally, the homogeneous trimeric assembly of all the variant SARS-CoV-2 S2P probes was confirmed by negative-stain electron microscopy (EM) (Fig 4). Although B.1.1.529 S2P probe appeared different in shape from the other S2P probes under negative-stain EM at pH 7, it exhibited mostly normal trimeric particle shape at pH 5.5 (S6 Fig), and all other characterization indicated B.1.1.529 S2P probe to be intact and well-folded, and to function as expected. It is unclear why at pH 7 this probe appeared different under negative-stain EM; it could possibly indicate some flexibility of the S2 subunit of B.1.1.529. Altogether, we succeeded in making biotinylated SARS-CoV-2 spike glycoproteins for variants of concern, including B.1.1.7. (Alpha), B.1.35s1 (Beta), P.1 (Gamma), B.1.617.2 and AY.1 (both are categorized under Delta), B.1.1.529 (Omicron), and several other notable variants.

**Fig 4.**
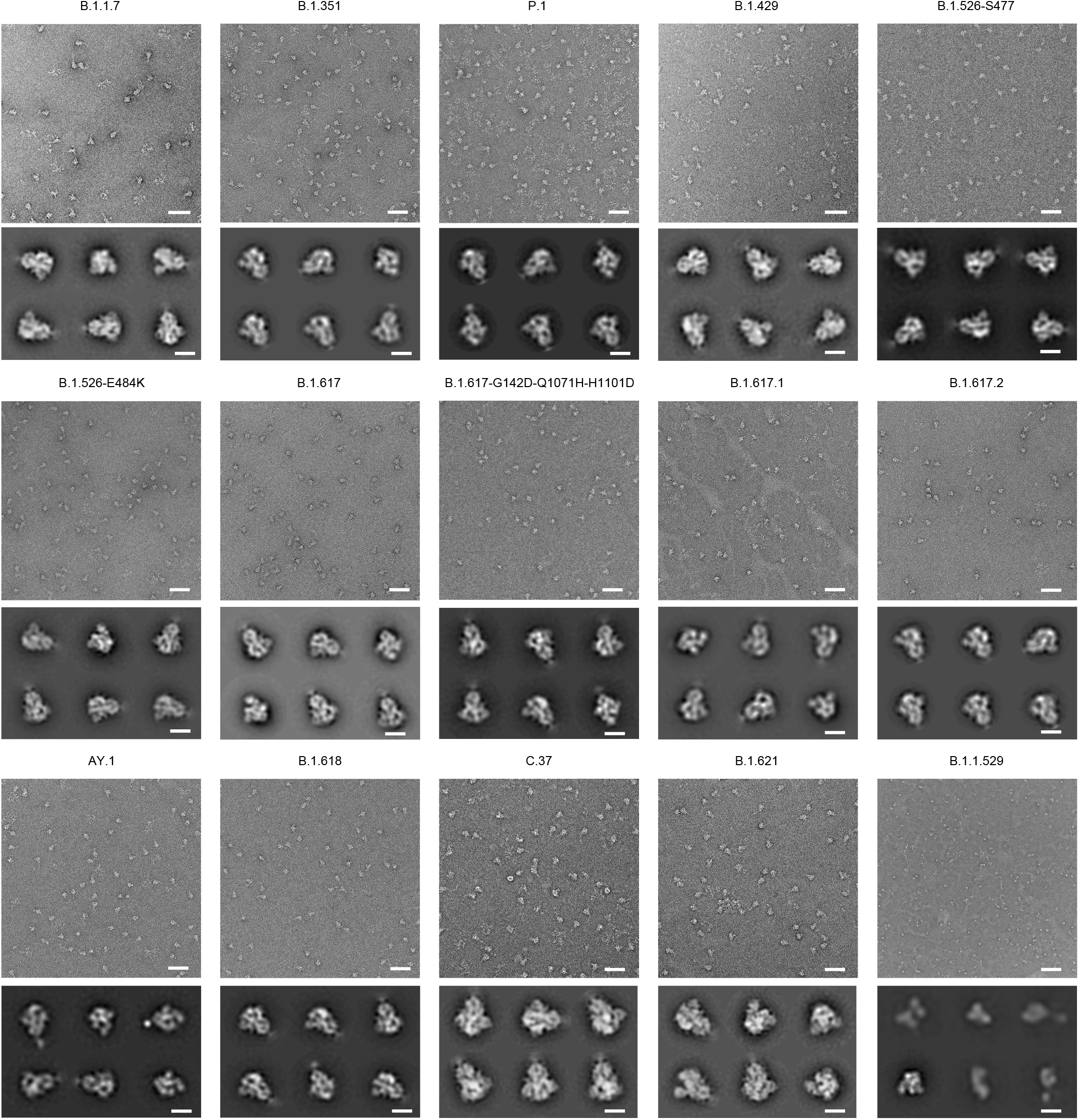
Negative stain EM of the biotinylated SARS-CoV-2 variant S2P probes shows individual trimeric spike to be well folded. The variants are labeled above each set of micrographs. The top panel of each set is the representative micrograph; the bottom panel shows the 2D-class averages. Scale bars in micrographs: 50 nm. Scale bars in 2D class average images: 10 nm. The B.1.1.529 S2P probe showed mostly trimeric particles, but appeared missing the S2 subunits at pH 7.5; however, at pH 5.5, it exhibited normal trimeric structure as other S2P probes (see S6 Fig).

### Molecular probes of variant SARS-CoV-2 NTD, RBD, RBD-SD1 regions

Recent reports of SARS-CoV-2 antibodies have shown that the receptor binding domain (RBD) and the N-terminal domain (NTD) are immunodominant and contain critical epitopes for neutralizing antibodies [34–38]. Therefore, the domain probes for variants are highly desirable for readout of immunogenicity and for isolation and characterization of neutralizing antibodies. Accordingly, we created separate molecular probes comprising the NTD (residues 14-305), the RBD (residues 329-526 for B.1.1.529 and residues 324-526 for other variants), and RBD with subdomain 1 (RBD-SD1, residues 319-591) with corresponding mutations listed in Fig 2. We designed and prepared 14 NTD probes, 12 RBD probes, and 12 RBD-SD1 probes from various variant sequences alongside their original WA-1 strain counterpart. The expression and purification were similar to those for S2P probes as described above. The overall yields were between 1.5-5.9 mg/L for the NTD probes, 3.7-11.3 mg/L for the RBD probes, and 3.8-18.3 mg/L for the RBD-SD1 probes, except for B.1.1.529 variant, which yielded 16.6 mg/L for NTD, mg/L for RBD, and 0.88 mg/L for RBD-SD1.

All purified biotinylated NTD probes ran as a single peak on size-exclusion chromatography (Fig 5A) and appeared as a single major band on SDS-PAGE (Fig 5B). The NTD bands look diffused and the observed molecular weights are higher than the theoretical value (∼36kDa) because the NTD is heavily *N*-link glycosylated. Biotinylation of the probes and their binding to the antibody panel were evaluated by BLI. All purified NTD probes bound well to streptavidin biosensors (Fig 5C). As expected, none of the NTD probes reacted to S2-directed or RBD-directed antibodies (Fig 5D). Unlike the moderate signals observed with the variant S2P probes, all the variant NTD probes clearly showed high responses to antibody S652-118, indicating that the epitope of S652-118 was conserved among SARS-CoV, WA-1 SARS-CoV-2, and all the SARS-CoV-2 variants investigated here. The relatively low reactivity of S652-118 to the S2P trimer probes was likely due to the restricted access to the S652-118 epitope by other domains in the trimer. Among the other three SARS-CoV-2 NTD-directed antibodies, 5-7 could bind to most of the NTD probes, indicating its epitope is unaffected by mutations in the NTD region of most of the variants. As observed with S2P probes, only NTD probes from B.1.617.2 and AY.1 showed low BLI responses to antibody 5-7. On the other hand, antibodies 2-51 and 5-24 showed a discerning difference in binding NTD probes in comparison with S2P probes. In addition to the variants that 2-51 and 5-24 could not recognize in the S2P probes, these two antibodies could not also recognize the variant B.1.429-W152C NTD probe (Fig 5D).

**Fig 5.**
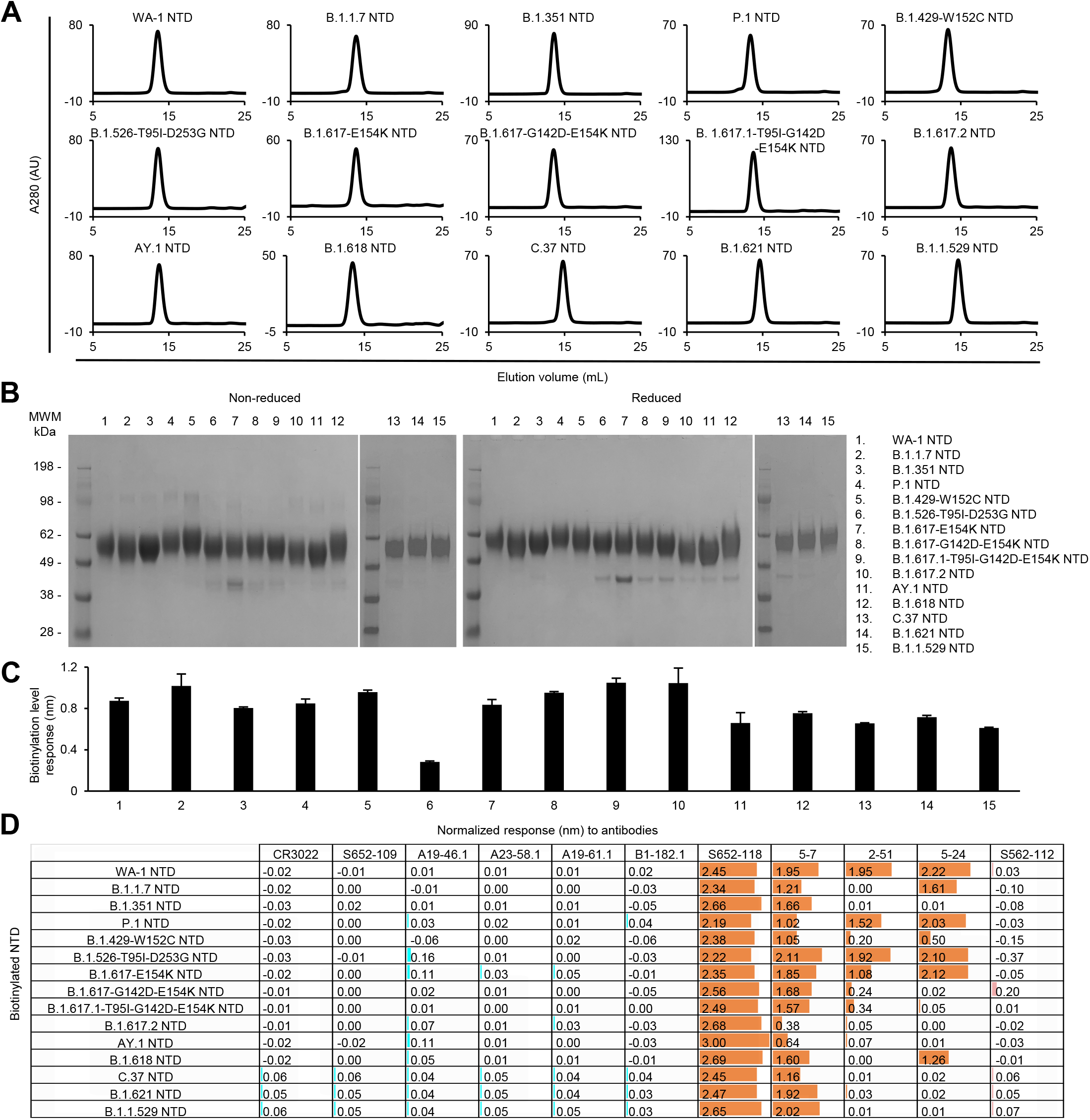
Characterization of biotinylated SARS-CoV-2 variant NTD probes confirm their homogeneity and antibody-binding specificity. (A) Size exclusion chromatography of purified biotinylated SARS-CoV-2 NTD probes, including original WA-1 NTD and 14 other variant NTD in PBS buffer. (B) SDS-PAGE of SARS-CoV-2 NTD variant probes with and without reducing agent. Molecular weight marker (MWM) and the WA-1 NTD probe were run alongside the variant NTD probes. (C) Biotinylation of the SARS-CoV-2 variant NTD probes. The level of biotinylation was evaluated by capture of NTD probes at 5 μg/ml onto streptavidin biosensors using Octet. Error bars represent standard deviation of triplicate measurements. (D) Antigenicity assessment of the SARS-CoV-2 variant NTD probes. Responses to RBD-directed, NTD-directed, and S2 subunit-directed antibodies were shown in cyan, orange, and rose, respectively. Bar scale between 0 and 3. Negative values not depicted.

For the RBD and RBD-SD1 regions, we produced 12 variant probes each along with the WA-1 RBD and RBD-SD1, because the B.1.526-E484K is the same as B.1.618, and the two B.1.617 variants are the same as the Kappa variant, B.1.617.1, in their RBD and SD1 sequences. All purified RBD and RBD-SD1 molecular probes appeared as a single peak on size-exclusion chromatography (Figs 6A and 7A) and ran as a single band on non-reducing and reducing SDS-PAGE (Figs 6B and 7B). The RBD and RBD-SD1 probes bound well to streptavidin biosensors (Figs 6C and 7C), and almost all showed high responses to every RBD-antibodies tested, except antibodies A19-46.1 and A19-61.1 (Figs 6D and 7D). A19-46.1 could not bind probes with L452R/Q mutation, as seen with S2P probes (Figs 3D, 6D and 7D). On the other hand, B.1.1.529 RBD only lost binding to the class 3 RBD antibody A19-61.1, unlike B.1.1.529 S2P, which could not recognize A19-61.1 nor the class 1 RBD antibodies A23-58.1 and B1-182.1. Overall, the binding results from variant subdomain probes provide molecular insights into antibody recognition of SARS-CoV-2 variants.

**Fig 6.**
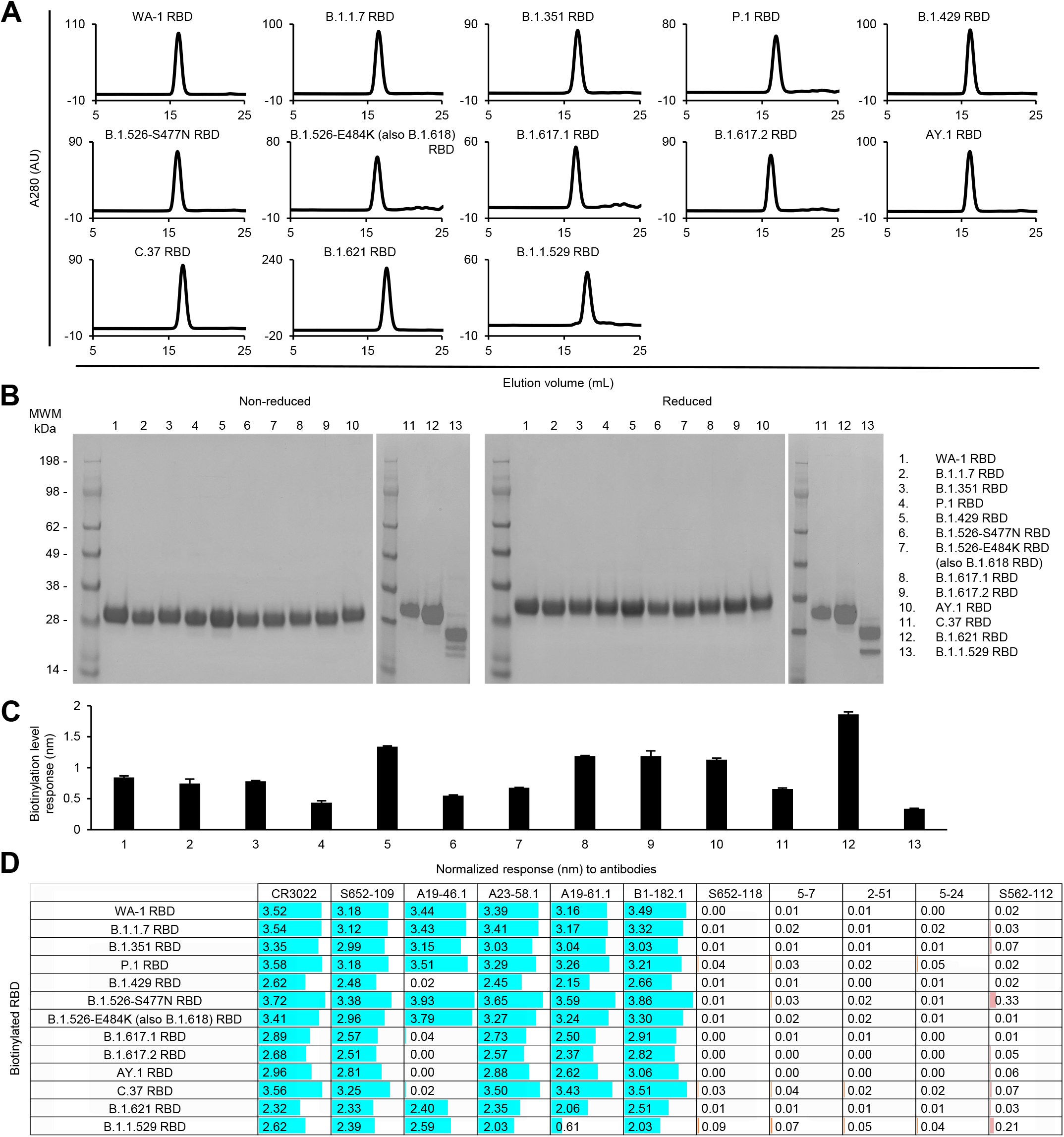
Characterization of biotinylated SARS-CoV-2 variant RBD probes confirm their homogeneity and antibody-binding specificity. (A) Size exclusion chromatography of the purified biotinylated SARS-CoV-2 RBD probes, including WA-1 RBD and 12 other variant RBD probes, in PBS buffer. (B) SDS-PAGE of SARS-CoV-2 RBD variant probes with and without reducing agent. Molecular weight marker (MWM) and the WA-1 RBD probe were run alongside the variant RBD probes. (C) Biotinylation of the SARS-CoV-2 variant RBD probes. The level of biotinylation was evaluated by capture of the RBD probes at 2.5 μg/ml onto the streptavidin biosensors using Octet. Error bars represent standard deviation of triplicate measurements. (D) Antigenicity assessment of the SARS-CoV-2 variant RBD probes. Responses to RBD-directed, NTD-directed, and S2 subunit-directed antibodies were shown in cyan, orange, and rose, respectively. Bar scale between 0 and 4. Variants with the L452R mutation did not binding antibody A19-46.1.

**Fig 7.**
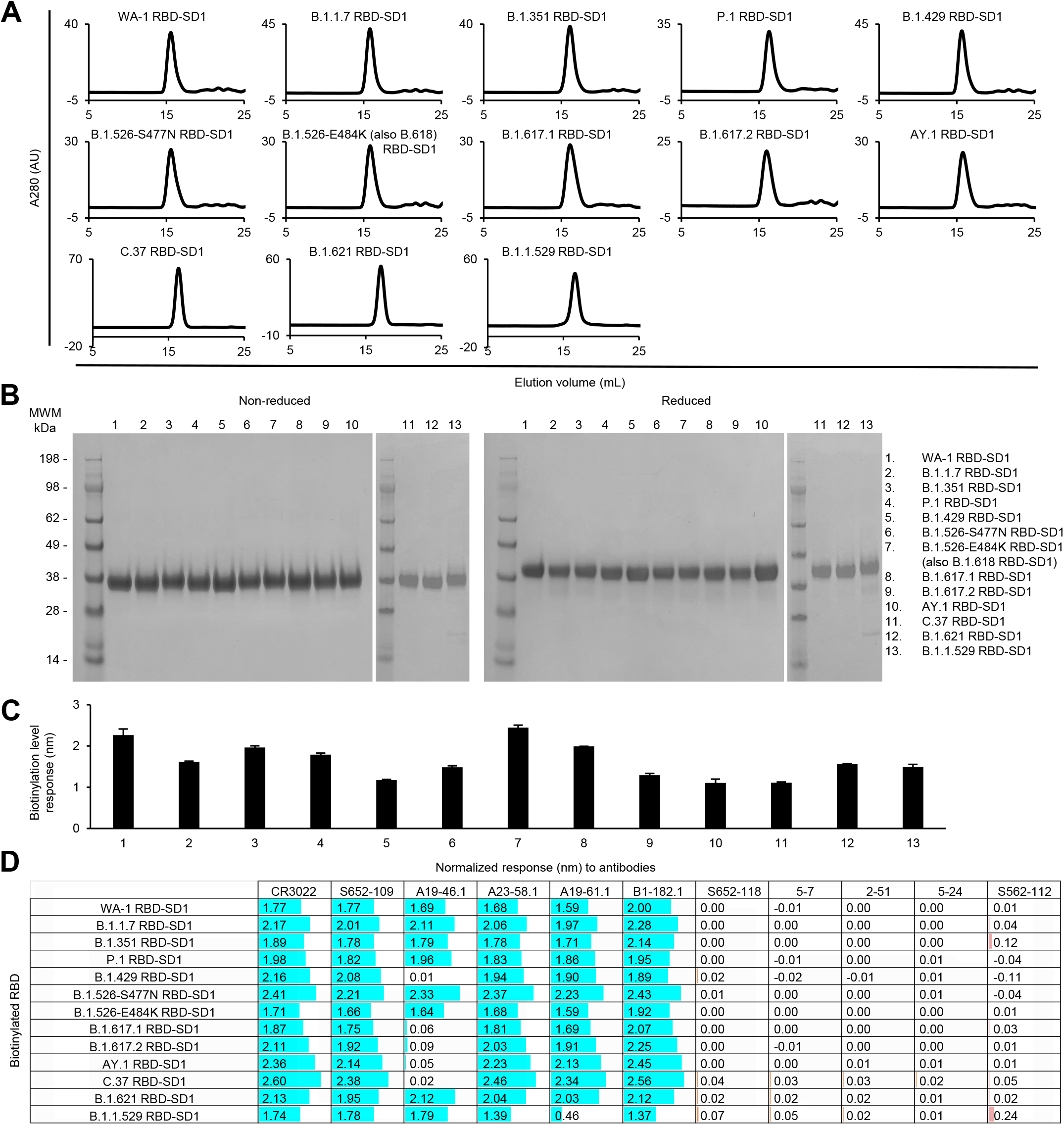
Characterization of biotinylated SARS-CoV-2 variant RBD-SD1 confirm their homogeneity and antibody-binding specificity. (A) Size exclusion chromatography of the purified biotinylated SARS-CoV-2 RBD-SD1 probes, including original WA-1 RBD-SD1 and 12 other variant RBD-SD1 in PBS buffer. (B) SDS-PAGE of SARS-CoV-2 RBD-SD1 variant probes with and without reducing agent. Molecular weight marker (MWM) and the WA-1 RBD-SD1 probe were run alongside the variant RBD-SD1 probes. (C) Biotinylation of the SARS-CoV-2 variant RBD-SD1 probes. The level of biotinylation was evaluated by capture of the RBD-SD1 probes at 5 μg/ml onto the streptavidin biosensors using Octet. Error bars represent standard deviation of triplicate measurements. (D) Antigenicity assessment of the SARS-CoV-2 variant RBD-SD1 probes. Responses to RBD-directed, NTD-directed, and S2 subunit-directed antibodies were shown in cyan, orange, and rose, respectively. Bar scale between 0 and 3. Negative values not depicted. Variants with the L452R mutation lost binding to A19-46.1.

### Binding analysis to yeast displaying monoclonal antibodies and convalescent immune libraries

To evaluate the variant molecular probes for their utilities in immunogenic assays and cell sorting, we used flow cytometry to measure probe binding to yeast displaying either antigen-specific antibodies or libraries cloned from B cells isolated from COVID-19 convalescent patients [39]. The biotinylated SARS-CoV-2 S2P, NTD, and RBD probes were freshly labeled with fluorochromes and used to sort the yeast cells displaying Fabs of S652-112, S652-118, and S652-109, antibodies isolated from a SARS-CoV-1 convalescent subject that bind SARS-CoV-2 S2, NTD and RBD, respectively [19], and Fabs of LY-555, CB6, REGN10933, REGN10987, A19-46.1, and A23-58.1 isolated from COVID-19 convalescent patients [18, 40, 41]. For S652-112 expressing yeast, S2P variant probes showed less than 2-fold (ranged 59%-167%) difference in binding relative to WA-1 S2P probe (Fig 8A and S2 Fig). The variant with the lowest binding had mutation A701V (B.1.351, 59%) in the S2 stem (Figs 1 and 7A). While more work is needed to define the S652-112 epitope, this data suggests it may reside near residue 701 in the S2 stem region (Fig 1). The NTD-reactive S652-118 expressing yeast bound to both S2P and NTD probes as expected (Fig 8A; S2 and S3 Figs). We found that S2P binding was 59-167% and NTD was 48-200% of WA-1 binding. We noted that S652-118 had reduced binding to B.1.429 S2P trimer (59%) but increased binding (146%) to NTD probe of the same variant. While these differences are subtle, they suggest that the decrease in S2P binding may not simply be due to the W152C substitution, which is located on a surface loop equally accessible between S2P and NTD probes, but rather may be due to the different accessibility of the epitope in the trimer versus the isolated NTD (Figs 2 and 8A; S2 and S3 Figs). Conversely, S652-118 binding to the probes of B.1.526 and AY.1 had opposite effects, with NTD being worse (48% and 48%, respectively) and slightly higher in the context of these S2P trimer variants (134% and 167%, respectively) (Fig 8A). This suggests that the S652-118 either has stabilizing contacts with other non-NTD domains or that NTD epitope is stabilized in context of the trimeric S-2P.

**Fig 8.**
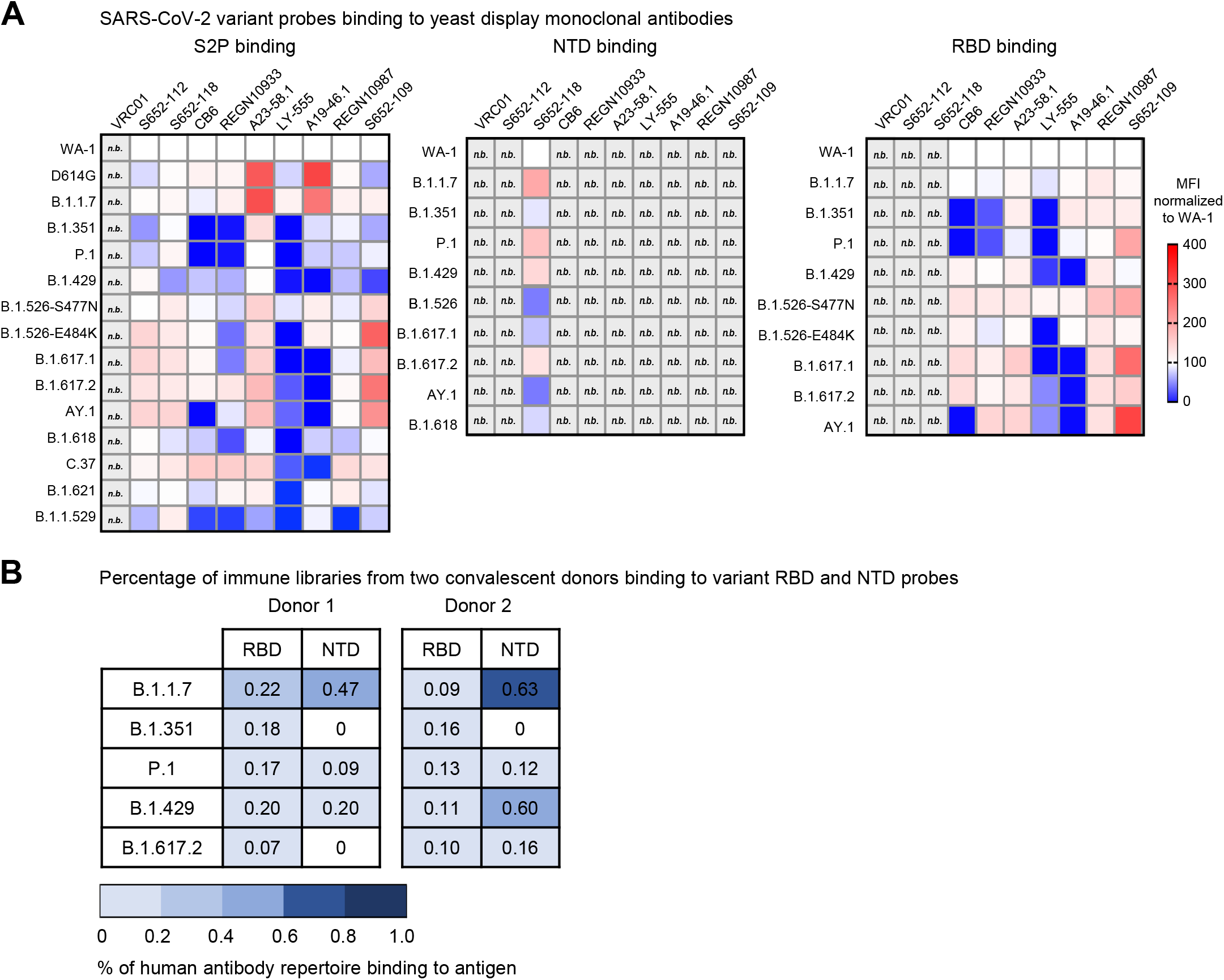
Interrogation of SARS-CoV-2 probes using yeast displaying anti-SARS-CoV-2 spike Fabs and immune libraries. (A) Binding of yeast expressing SARS-CoV cross-reactive Fabs (S652-112, S652-118, and S652-109), SARS-CoV-2 Fabs (CB6, REGN10933, A23-58.1, LY-555, A19-46.1, and REGN10987) or HIV targeting VRC01 Fab to SARS-CoV-2 WA-1, variant D614G, B.1.1.7, B.1.351, P.1, B.1.429, B.1.526-S477N, B.1.526-E484K, B.1.617.1, B.1.617.2, AY.1, B.1.618, C.37, B.1.621, and B.1.1.529 spike ectodomain S2P-APC (left), NTD-BV711 (middle), and RBD-BV421 (right). Binding to the indicated yeast displayed antibody was measured with flow cytometry. Data are shown as Mean Fluorescence Intensity (MFI) for the same antibody against the parental strain WA-1 probe. The normalized percent change is indicated by a color gradient from red (increased binding, Max 400%) to white (no change, 100%) to blue (complete loss of binding, 0%). Within each class of probe (i.e., S2P, NTD and RBD), yeast expressing Fab but do not bind any probes are shown in light grey and marked as “n.b.”. (B) Percentage of human antibody repertoire binding to NTD and RBD of SARS-CoV-2 variants. Natively paired antibody heavy and light chains were captured from COVID-19 convalescent patient immune repertoires and displayed as Fabs on the surface of yeast. Probe binding to the yeast-displayed Fabs was analyzed against pre-conjugated SARS-CoV-2 biotinylated variant probes and measured with flow cytometry. Numbers represent the percentage of binding to the biotinylated probes after subtracting the background fluorescence.

We next evaluated the binding of S2P and RBD probes to yeast expressing RBD Class I (e.g., CB6, REGN10933, A23-58.1), Class II (e.g., LY-555, A19-46.1) or Class III (e.g., REGN10987, S652-109) antibody Fab regions. We noted that relative to WA-1, binding of D614G S2P and B.1.1.7 S2P probes to yeast expressing A23-58.1 and A19-46.1 was increased 260-275% and 292-214% respectively (Fig 8A and S2 Fig). A23-58.1 is a Class I antibody that binds to the tip of RBD when it is in the up position and A19-46.1 is a Class II antibody that binds to RBD when it is in the up or down position. Neither antibody shows increased binding to the B.1.1.7 RBD probe (Fig 7A; S2 and S4 Figs). This indicates that the increased binding is not likely due to the presence of the N501Y mutation in B.1.1.7 but rather reflects a preference for these antibodies to bind RBD up and an increased probability of RBD being present in the up position. In addition to and consistent with previously published results, binding of A23-58.1 yeast was not decreased to any of the S2P or RBD variant probes, and A19-46.1 yeast showed decreased binding to variant S2P and RBD probes containing L452R mutations. In line with previous reports showing significant loss of binding and neutralization by CB6 and REGN10933 to B.1.351 and P.1, yeast bearing CB6 and REGN10933 showed decreased binding to S2P and RBD (Fig 8A; S2 and S4 Figs). Similarly, S2P probes with mutations at E484 also decreased binding to REGN10933 yeast that was not present in the context of RBD monomer probe. Notably, for the Class II antibody LY-555, binding was reduced in the presence of E484Q/K in both S2P and RBD trimer. Taken together these data validate the use of the biotinylated S2P, NTD and RBD probes for immunogenic assays and cell sorting.

Next, we used RBD and NTD probes from five variants, including B.1.1.7, B.1.351, P.1, B.1.429, and B.1.617.2, to cross examine human antibody repertoire from two convalescent donors that were infected with WA-1 strain. The natively paired VH:VL antibody genes from B cells isolated from PBMCs were cloned into a yeast display vector to generate a yeast library from each donor, expressing human antibody Fabs on yeast surface. The sorting results showed that both libraries responded to all five variant RBD probes with percentage of antigen-positive ranged between 0.07-0.22 % for one library and 0.09-0.16 % for the other (Fig 8B). This result is consistent with the understanding that prior exposure to SARS-CoV-2 or inoculation with vaccines designed based on WA-1 strain still protects against these variants. However, when sorting with variant NTD probes, donor 1 library did not bind B.1.351 or B.1.617.2, and donor 2 library bound poorly to B.1.351, indicating that the NTD-directed antibodies elicited by natural infection are susceptible to mutations in NTD of B.1.351 and B.1.617.2.

Overall, these results validate the utilities of the variant probes in identification and characterization of antibodies against SARS-CoV-2 variants of concern and in assessment of immune responses and serum antibody specificity.

## Discussion

Our initial interest in developing molecular probes comprising key targets of the SARS-CoV-2 spike related to their use in defining and monitoring elicited responses and in facilitating antibody identification and characterization. Other uses in diagnostics (to assess sera reactivity and to provide sensitive markers of infection) and in pathogenesis (to delineate susceptible cells that virus might infect) were also possible. In these uses, one utility of the probes was in delineating the region of the spike recognized by antibodies, e.g., NTD, RBD, SD1 or elsewhere on the ectodomain, and when used in this way, the probes defining RBD and RBD-SD1 were somewhat redundant, as there is considerable overlap between these two probe regions. The poor expression of Omicron RBD (0.18 mg/L) versus the higher expression of Omicron RBD-SD1 (0.88 mg/L), however, demonstrates an additional utility of the RBD-SD1 construct, its increased expression.

The emergence of SARS-CoV-2 variants highlights an additional purpose, the tracking and characterization of the recognition by sera and antibodies of viral variants. Here, we have developed a matrix of probe regions (spike ectodomain, NTD, RBD, and RBD-SD1) and their variants (Alpha through Omicron) and demonstrated the utilities of these probes in analysis of sera and antibodies. It will be interesting to see how the field utilizes this matrix of probes – individually, such as with the Omicron spike or RBD, as a series, such as assessing sera with RBDs from Alpha through Omicron, or as a matrix. To facilitate usage of these probes in COVID-19 related research, the full matrix of probe regions and variants is being made available at Addgene for dissemination.

## Materials and methods

### Cell lines

Expi293F and FreeStyle 293-F cells were purchased from Thermo Fisher Scientific. The cells were maintained following manufacturer’s suggestions and were used as described in detail below.

### Construction of expression plasmids for wildtype and variant SARS-CoV-2 molecular probes

DNA sequences encoding the wildtype or variant SARS-CoV-2 spike or specific domains were cloned into a pVRC8400-based expression vector by GeneImmune. The gene of interest was inserted between the DNA sequences coding for a N-terminus purification component and a C-terminus biotinylation tag. The N-terminus purification component was composed of a single chain human Fc with knob-in-hole mutations, a ‘GGSGGGGSGG’ linker, and an HRV3C protease cleavage site (scFc3C) for purification with protein A resin and tag cleavage. And the C-terminus comprised a ‘GGGLVPQQSG’ 10lnQQ linker followed by an AVI tag for biotinylation. For the S2P probe constructs, the insertion included the wildtype spike protein residues 14 to 1208, or the corresponding spike gene from variants, with the S1/S2 RRAR furin cleavage site replaced by GSAS, the two proline stabilization mutations K986P and V987P, a GSG linker, and the T4-phage fibritin trimerization domain (foldon) as described by Wrapp and colleagues [28]. For the NTD probe constructs, spike gene coding for residues 14-305 was cloned. For the RBD probe constructs, spike gene coding for residues 324-526 was cloned. For the RBD-SD1 probe constructs, spike gene coding for residues 319-591 was cloned. Plasmids for the wildtype and variant SARS-CoV-2 proteins with AVI tag generated in this study have been deposited with Addgene (www.addgene.org) with accession numbers listed in S1 Table.

### Expression and preparation of wildtype and variant SARS-CoV-2 molecular probes

The wildtype and variant SARS-CoV-2 molecular probes were produced by transient transfection of FreeStyle 293-F cells [28] and in-process biotinylation [19] as previously described. Briefly, 1 mg of the plasmid encoding the scFc3C tagged and AVI tagged SARS-CoV-2 target protein and 3 ml of the Turbo293 transfection reagent (Speed BioSystems), each in 20 ml Opti-MEM (Thermo Fisher Scientific), were pre-mixed and transfected into FreeStyle 293-F cells at 1 mg plasmid per 0.8 L cells (2×10^6^ cells/ml). Cells were incubated in shaker at 120 rpm, 37 °C, 9% CO2. The next day following transfection, 80 ml HyClone SFM4HEK293 medium (Cytiva) and 80 ml FreeStyle 293 Expression Medium (Thermo Fisher Scientific) were added to each 0.8 liter of cells. The transfected cells were allowed to grow for 6-7 days in total before the supernatant was harvested by centrifugation and filtration. Next, the supernatant was incubated with 5ml of PBS-equilibrated protein A resin for two hours, after which the resin was collected and washed with PBS. The captured SARS-CoV-2 protein was then biotinylated using the BIRA500 kit (∼2.5 μg per 10 nmol AVI substrate) (Avidity) and cleaved from the Fc purification tag concurrently with 200 μg of HRV3C prepared as described [42]. After incubation at 4°C overnight, the liberated protein was collected, concentrated, and applied to a Superdex 200 16/600 gel filtration column equilibrated with PBS. Peak fractions were pooled and concentrated to 1 mg/ml. Finally, probes were run on SDS-PAGE to examine molecular size and purity.

### Expression and preparation of the ACE2 receptor

The human ACE2 (1-740aa) expression plasmid was constructed with a monomeric human Fc tag and an 8xHisTag at the 3’-end of the ACE2 gene. After transient transfection of Expi293F cells, the cell culture supernatant was harvested 5 days post transfection and loaded onto a protein A affinity column. The Fc-tagged protein was eluted with IgG elution buffer and then dialyzed against PBS.

### Expression and preparation of antibodies

DNA sequences of heavy and light chain variable regions of antibody CR3022 [32] and of donor S652 antibodies, S652-109, S652-118, and S652-112, and of SARS-CoV-2 antibodies A19-46.1, A23-58.1, A19-61.1, B1-182.1 [18], 5-7 [33], 2-51, and 5-24 [34] were synthesized and subcloned into the pVRC8400 vector, as described previously [43]. For antibody expression, equal amounts of heavy and light chain plasmid DNA were transfected into Expi293F cells using Turbo293 transfection reagent (Speed BioSystems). The transfected cells were cultured in shaker incubator at 120 rpm, 37 °C, 9% CO2 for 5 days before the culture supernatants were harvested, filtered, and purified using on AmMag SA semiautomated system (Genscript) and AmMag Protein A beads (Genscript) or loaded on a protein A (Cytiva) column. After washing the column or beads with PBS, each antibody was eluted with an IgG elution buffer (Pierce) and immediately neutralized with one tenth volume of 1M Tris-HCl pH 8.0. Eluted antibodies were then buffer exchanged with PBS, pH 7.4, using 10,000 MWCO dialysis cartridges (Pierce) overnight and were confirmed by SDS-PAGE before use.

### Bio-Layer Interferometry

An Octet HTX instrument (Sartorius) was used to analyze biotinylation level and antigenicity of the molecular probes and the receptor recognition of the S2P probes. Assays were performed at 30°C in tilted black 384-well plates (Geiger Bio-One) in PBS with 1% BSA with agitation set to 1,000 rpm. Before running the assays, the streptavidin biosensors were equilibrated in PBS with 1% BSA for at least 20 minutes. Biotinylated SARS-CoV-2 S2P (10 μg/ml), NTD (5 μg/ml), RBD (2.5 μg/ml), and RBD-SD1 (5 μg/ml) were loaded on to streptavidin biosensors for 60 seconds. Binding to biotinylated probes was measured by dipping immobilized probes into solutions of ACE2 receptor at 500 nM or antibodies at 200 nM for 180 seconds. Measurements were assessed in triplicate. Parallel correction to subtract systematic baseline drift was carried out by subtracting the measurements recorded for a loaded sensor dipped into buffer only control well. The response values at the end of association step were reported.

### Negative-stain electron microscopy

Purified SARS-CoV-2 S2P variant probes were diluted to approximately 0.02 mg/ml with buffer containing 10 mM HEPES, pH 7, and 150 mM NaCl. A 4.7-µl drop of the diluted sample was applied to a glow-discharged carbon-coated copper grid. The grid was washed three times with the same buffer, and adsorbed protein molecules were negatively stained by applying consecutively three drops of 0.75% uranyl formate. Micrographs were collected at a nominal magnification of 100,000x (pixel size: 0.22 nm) using SerialEM [44] on an FEI Tecnai T20 electron microscope operated at 200 kV and equipped with an Eagle CCD camera or at a nominal magnification of 57,000x (pixel size: 0.25 nm) on a Thermo Scientific Talos F200C electron microscope operated at 200 kV and equipped with a Ceta camera. Particles were picked automatically using in-house written software (Y.T., unpublished). Reference-free 2D classification was performed with Relion 1.4 [45].

### Probe conjugation

Biotinylated SARS-CoV-2 proteins were conjugated using either allophycocyanin (APC) streptavidin (full-length spike proteins, every monomer labeled), Brilliant Violet 421 (BV421) streptavidin (RBD proteins), or Brilliant Violet 711 (BV711) streptavidin (NTD proteins). Reactions were prepared at a 4:1 molecular ratio of protein to streptavidin. Labeled streptavidin was added in ⅕ increments, with incubations at 4°C (rotating) for 20 minutes in between each addition. When labelled, probes were titrated over monoclonal yeast display to determine an optimal staining concentration in the range of 10 to 0.1 ng per ul of staining solution.

### Analysis of probe binding to monoclonal yeast

Monoclonal yeast displays were created, expressed, and analyzed as previously published [39]. Briefly, VH and VL regions of VRC01 (as a negative control) [46], S652-118, S652-112, S652-109 [19], and SARS-Cov2-specific antibodies LY-555 [40], CB6 [47], REGN10933, REGN10987 [48], A19-46.1, and A23-58.1 [18] were codon optimized for yeast expression using JCat [49], synthesized and cloned by Genscript into pCT-VHVL-K1 or pCT-VHVL-L1 yeast expression vectors [39]. Yeast display vectors were linearized and Saccharomyces cerevisiae strain AWY101 (MATα AGA1::GAL1-AGA1::URA3 PDI1::GAPDH-PDI1::LEU2 ura3–52 trp1 leu2Δ1 his3Δ200 pep4::HIS3 prb1Δ1.6R can1 GAL) was transformed. Yeast cells were maintained in YPD medium (20 g/L dextrose, 20 g/L peptone, and 10 g/L yeast extract). After yeast transformation, cells were routinely maintained in SDCAA selection medium (20 g/L dextrose, 6.7 g/L yeast nitrogen base, 5 g/L casamino acids, 8.56 g/L NaH_2_PO_4_·H_2_O, and 10.2 g/L Na_2_HPO_4_·7H_2_O). Fab display was induced by incubating yeast in SGDCAA induction medium (SDCAA with 20 g/L galactose, 2 g/L dextrose). Two days after induction, 1×10^6^ yeast cells were incubated in staining buffer (phosphate buffered saline + 0.5% BSA + 2mM EDTA) containing anti-Flag fluorescein isothiocyanate antibody (2 μg/mL; clone M2, Sigma-Aldrich) and the probes for 30 minutes at RT in darkness prior to washing 2 times in ice cold staining buffer. Fab expressing yeast (FLAG+) were analyzed for their capacity to bind to the indicated probes using a BD LSRFortessa X-50 Cell Analyzer (BD Biosciences). For chequerboard single probe stain experiments yeast were incubated with 1 ng/ul of S2P (APC), 1 ng/ul of RBD (BV421) or 1 ng/ul of NTD (BV711). Obtained flow cytometry data was further analyzed using FlowJo 10.6.1 software (BD Biosciences). The gating tree for antibody binding is shown in S1 Fig. Heat maps were created using GraphPad Prism 8 software (GraphPad Software Inc.).

### Analysis of probe binding to human antibody repertoires expressed in yeast

B cells were isolated from cryopreserved PBMCs of SARS-CoV-2 convalescent donors as previously described [16, 39, 50–52]. Briefly, B cells were enriched by CD27+ selection and stimulated *in vitro* for five days to enhance transcription of antibody genes. Single B cells were captured in emulsion droplets consisting of lysis buffer and oligo(dT)-coated magnetic beads for single-cell mRNA capture. Overlap extension RT-PCR was used to transform separate heavy (VH) and light (VL) variable genes into a single, physically linked VH:VL cDNA amplicons. cDNA amplicons encoding heavy and kappa light chain variable regions were cloned into a yeast display vector for expressing antibody VH:VL genes as Fabs for functional screening via FACS and NGS as previously reported [16, 39, 51, 52]. Briefly, AWY101 yeast expressing Fabs were cultured in SGCAA media (Teknova) with 2 g/L dextrose (SGDCAA) for 36 hrs at 20°C to induce Fab expression for surface display. In the first round of screening, 3×10^7^ Fab-expressing yeast cells were stained and washed with ice-cold staining buffer (phosphate-buffered saline (PBS) with 0.5% BSA and 2mM EDTA). Washed yeast libraries were stained in dark and at 4°C with biotinylated antigens pre-conjugated with SA-PE (Streptavidin, R-Phycoerythrin Conjugate, Thermo Fisher Scientific) or SA-APC (Streptavidin, Allophycocyanin Conjugate, Thermo Fisher Scientific), and an anti-FLAG fluorescein isothiocyanate (FITC) monoclonal antibody (clone M2-FITC, Sigma-Aldrich) was added as a marker for Fab expression. Sorting was performed on a SONY MA900 cell sorter and gates for sorting were drawn as previously described [39, 53]. Sorted yeast cells were collected and cultured in SDCAA (pH 4.5) for 12-24 h for subsequent rounds of enrichment for binding against antigens. Separate Fab expressing yeast cells were sorted for each library (referred as VL+) initially by staining yeast input libraries with anti-FLAG-FITC conjugated mAb.

## Author Contributions

Conceptualization: I-Ting Teng, Peter D. Kwong.

Data curation: I-Ting Teng, Alexandra F. Nazzari, Misook Choe, Matheus Oliveira de Souza, Yuliya Petrova, Yaroslav Tsybovsky, Tracy J. Ruckwardt, John Misasi.

Formal analysis: I-Ting Teng, Matheus Oliveira de Souza, Bharat Madan.

Funding acquisition: Brandon J. DeKosky, John R. Mascola, Nancy J. Sullivan, Peter D. Kwong. Investigation: I-Ting Teng, Alexandra F. Nazzari, Matheus Oliveira de Souza, Yuliya Petrova, Yaroslav Tsybovsky, Mykhaylo Artamonov, Aric Huang, Sheila N. Lopez Acevedo, Xiaoli Pan, Tracy J. Ruckwardt.

Methodology: I-Ting Teng, Tracy J. Ruckwardt, Brandon J. DeKosky, John Misasi, Tongqing Zhou, Peter D. Kwong.

Project administration: Peter D. Kwong.

Resources: Misook Choe, Tracy Liu, Baoshan Zhang.

Supervision: Peter D. Kwong.

Writing – original draft: I-Ting Teng, Shuishu Wang, Peter D. Kwong.

Writing – review & editing: I-Ting Teng, Shuishu Wang, Peter D. Kwong.

## Acknowledgements

We thank J. Stuckey for assistance with figures and members of the Vaccine Research Center for discussions or comments on the manuscript.

---

**S1 Fig.**
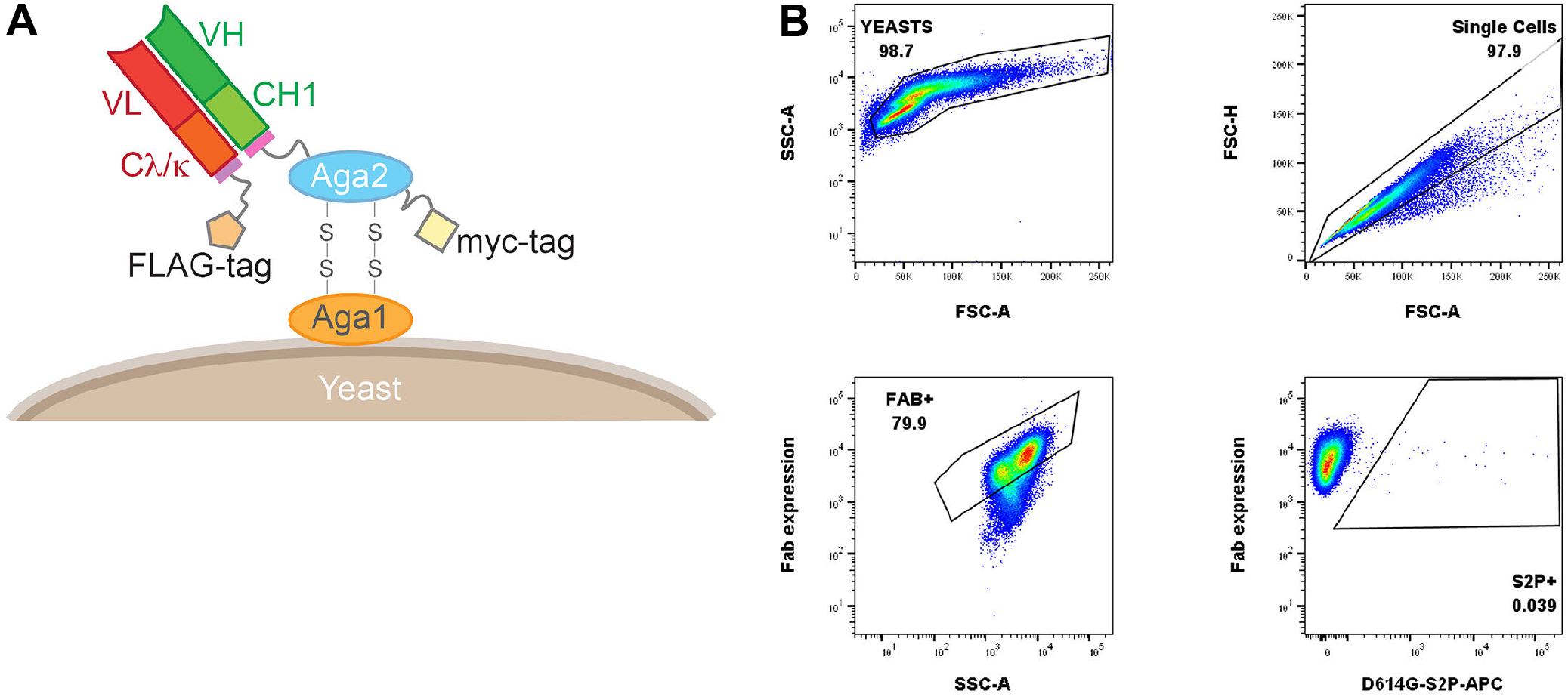
Yeast Fab display and gating tree for yeast display analysis of probe binding. (A) Saccharomyces cerevisiae strain AWY101 transfected with yeast display vector and Fab display is induced by incubating yeast in galactose containing media. The presence of Fab expressed on the yeast surface can be detected by staining with an anti-Flag antibody and analyzing using flow cytometry. (B) Induced yeast bearing Fabs of interested are analyzed by the indicated gating strategy. Singlets are analyzed for Fab expression and the proportion of probe binding determined within this population of yeast. Shown is a representative data.

**S2 Fig.**
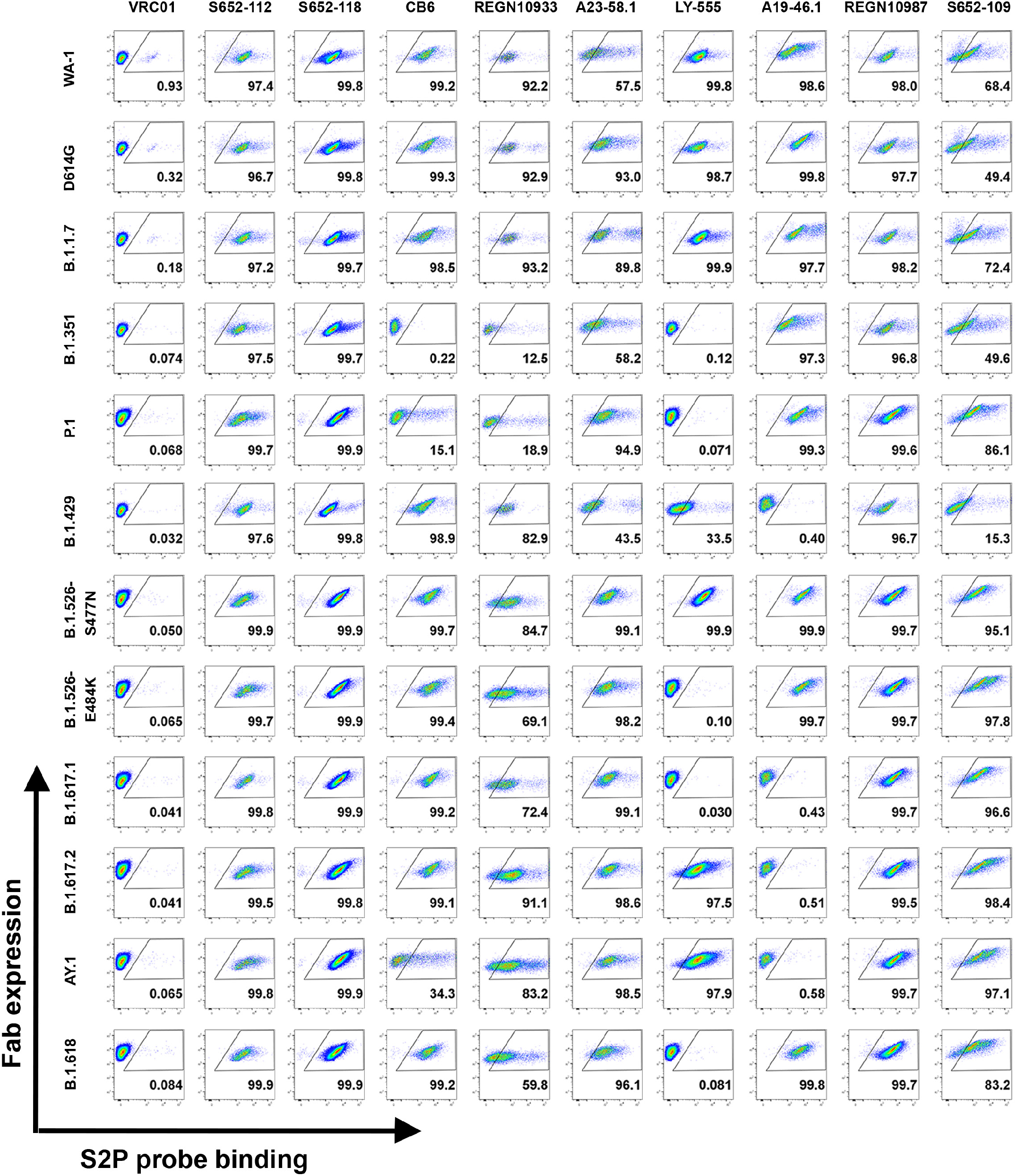
Yeast SARS-CoV cross-reactive and SARS-CoV-2 Fab binding to SARS-CoV-2 antigenic S2P probes. Binding of yeast expressing SARS-CoV cross-reactive Fabs (S652-118, S652-112, and S652-109), SARS-CoV-2 Fabs (LY-555, CB6, REGN10933, REGN10987, A19-46.1, and A23-58.1) or HIV targeting VRC01 Fab to SARS-CoV-2 VOC, VOI and other variant antigenic probes: WA-1, D614G, B.1.1.7, B.1.351, P.1, B.1.429, B.1.526-S477N, B.1.526-E484K, B.1.617.1, B.1.617.2, AY.1, and B.1.618 S2P (APC).

**S3 Fig.**
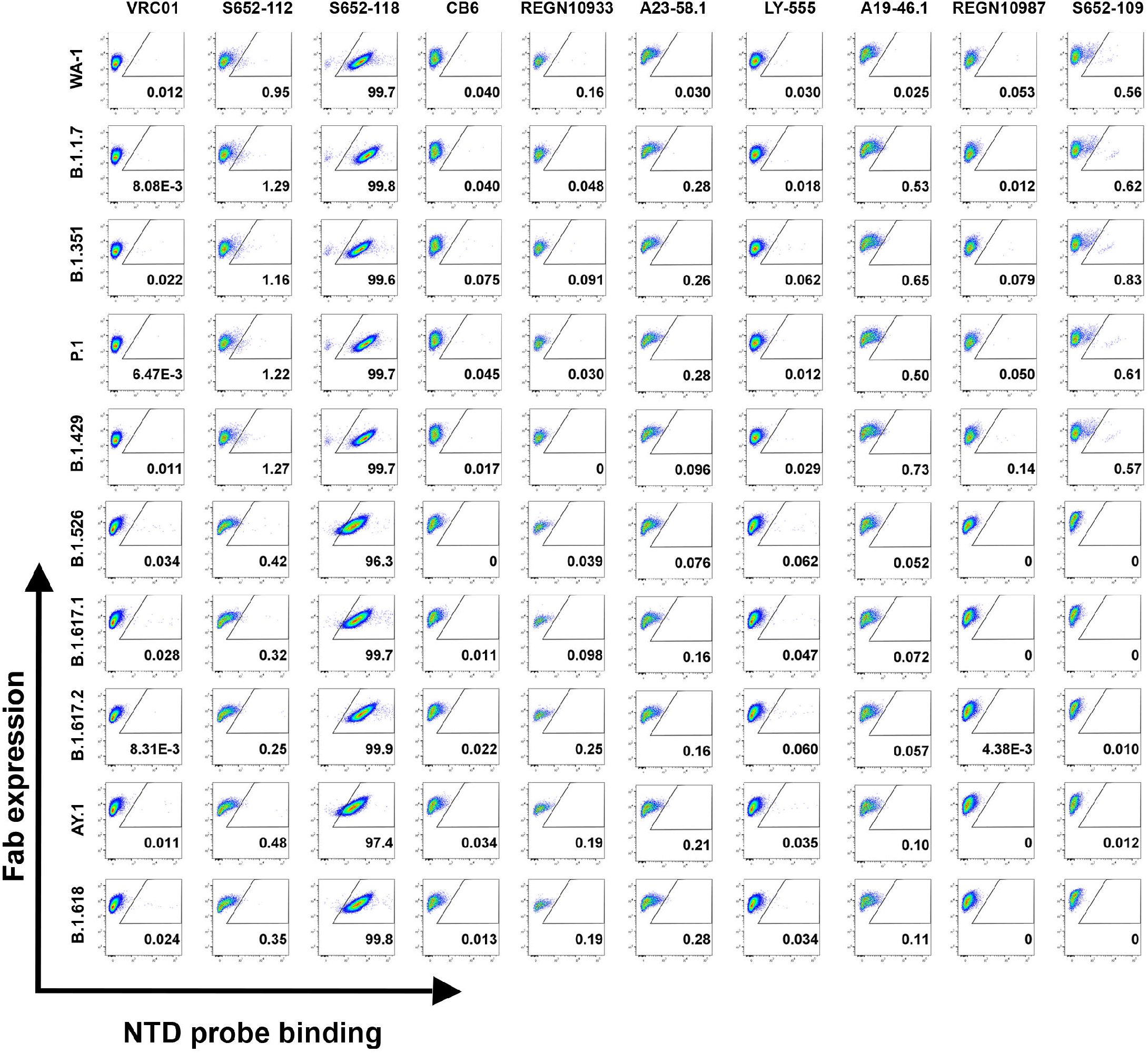
Yeast SARS-CoV cross-reactive and SARS-CoV-2 Fab binding to SARS-CoV-2 antigenic NTD probes. Binding of yeast expressing SARS-CoV cross-reactive Fabs (S652-118, S652-112, and S652-109), SARS-CoV-2 Fabs (LY-555, CB6, REGN10933, REGN10987, A19-46.1, and A23-58.1) or HIV targeting VRC01 Fab to SARS-CoV-2 VOC, VOI and other variant antigenic probes: WA-1, B.1.1.7, B.1.351, P.1, B.1.429, B.1.526, B.1.617.1, B.1.617.2, AY.1, and B.1.618 NTD (BV711).

**S4 Fig.**
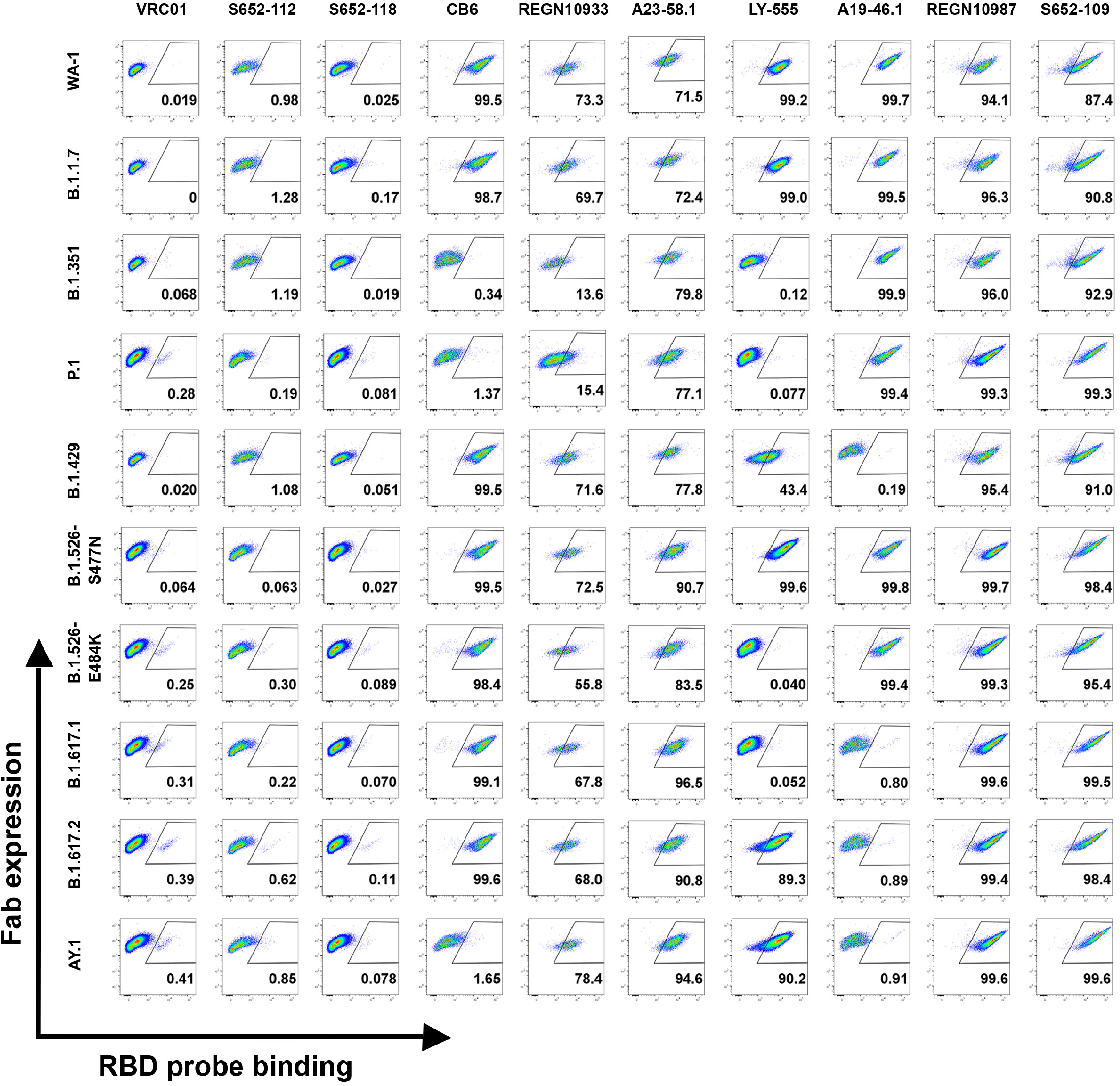
Yeast SARS-CoV cross-reactive and SARS-CoV-2 Fab binding to SARS-CoV-2 antigenic RBD probes. Binding of yeast expressing SARS-CoV cross-reactive Fabs (S652-118, S652-112, and S652-109), SARS-CoV-2 Fabs (LY-555, CB6, REGN10933, REGN10987, A19-46.1, and A23-58.1) or HIV targeting VRC01 Fab to SARS-CoV-2 VOC, VOI and other variant antigenic probes: WA-1, B.1.1.7, B.1.351, P.1, B.1.429, B.1.526-S477N, B.1.526-E484K, B.1.617.1, B.1.617.2, and AY.1 RBD (BV421).

**S5 Fig.**
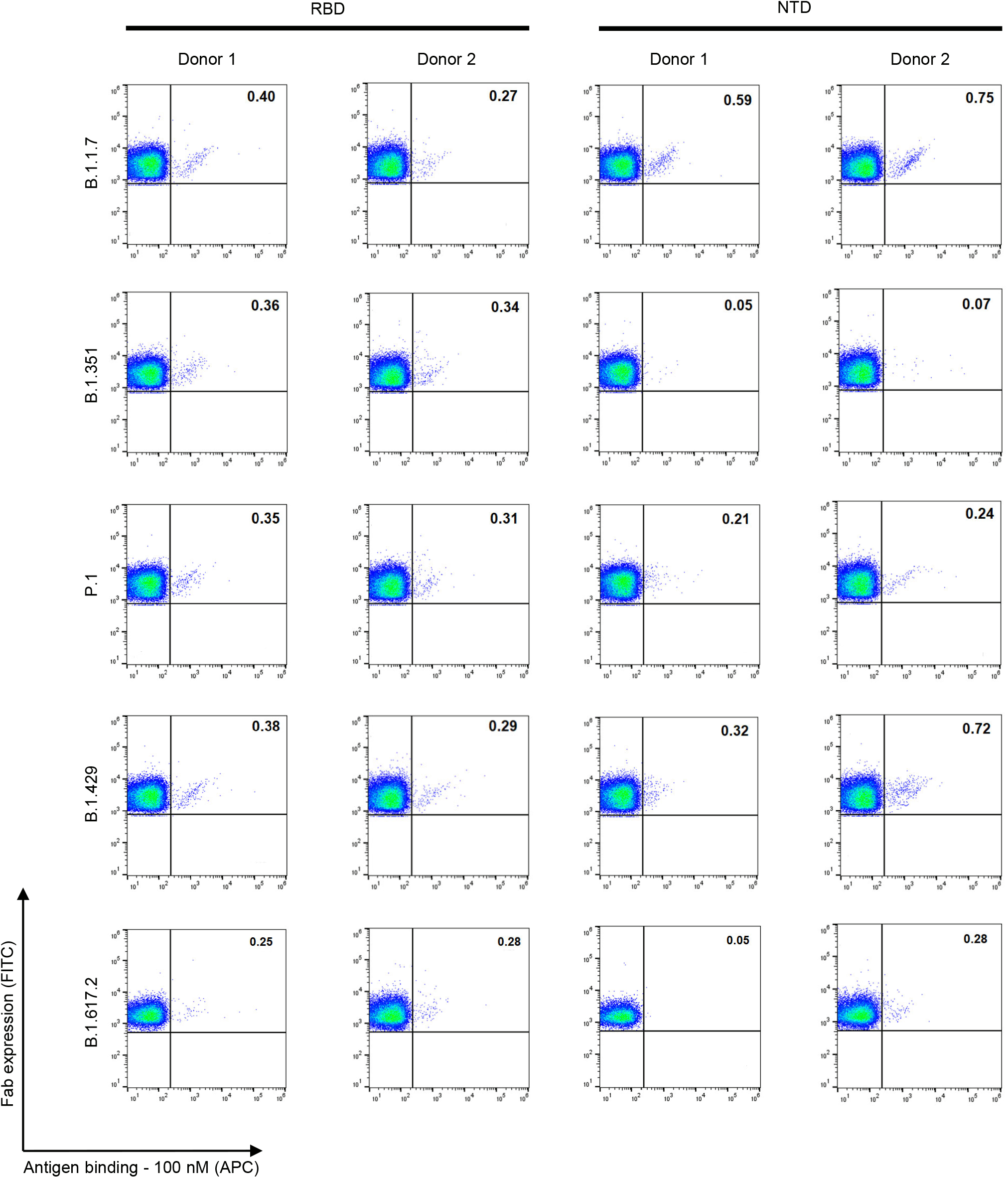
Yeast expressing human antibody repertoire binding to SARS-CoV-2 antigenic RBD and NTD probes. Binding of yeast expressing SARS-CoV-2 libraries (donor 1 and donor 2), targeting RBD and NTD of SARS-CoV-2 variants: B.1.1.7, B.1.351, P.1, B.1.429, and B.1.617.2

**S6 Fig.**
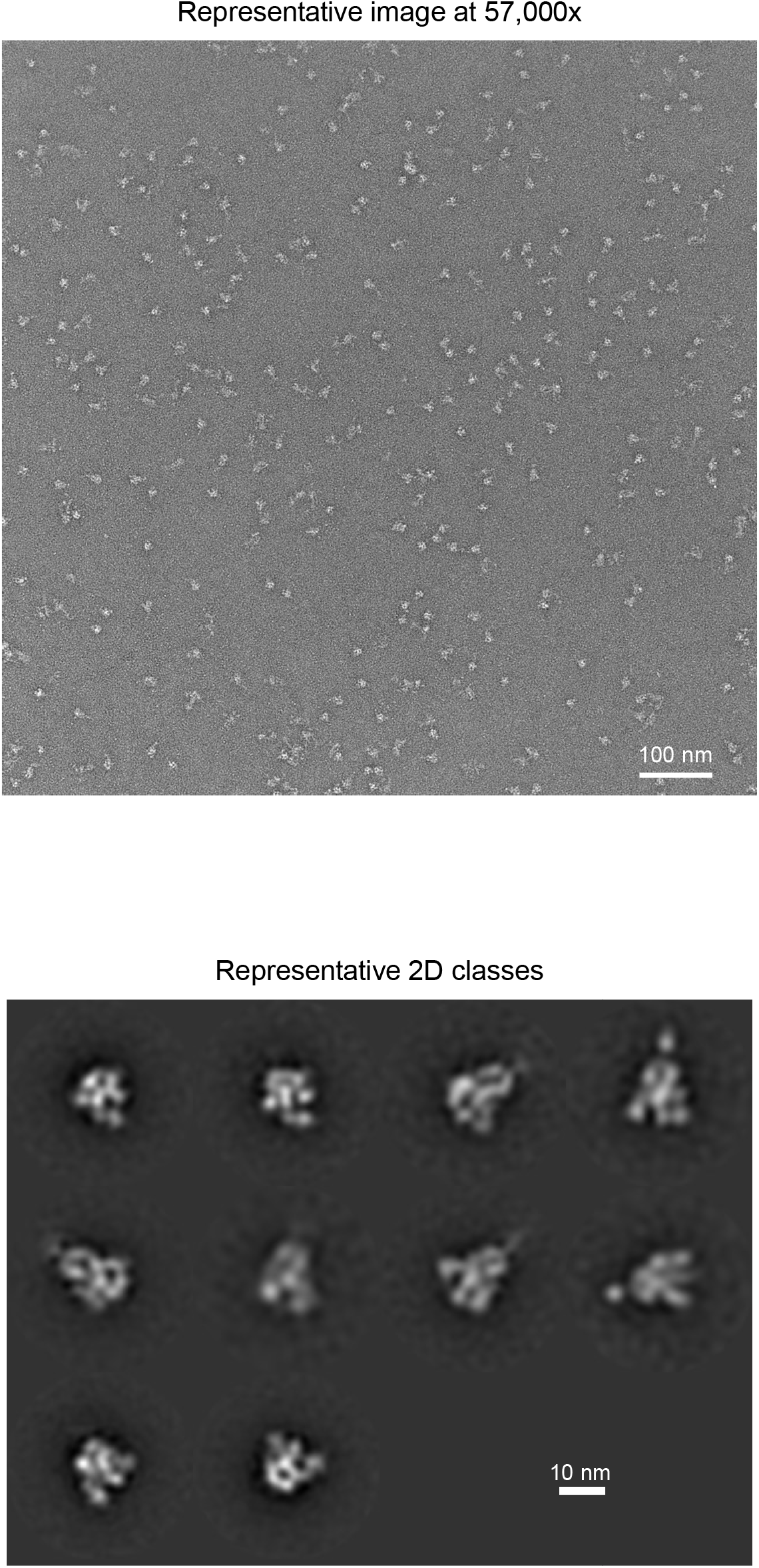
Negative-stain EM of the biotinylated SARS-CoV-2 Omicron variant S2P probes at pH 5.5 shows individual trimeric spike to be well folded. The top panel is the representative micrograph; the bottom panel shows the 2D-class averages. Sizes of scale bars are as indicated. At pH 5.5, B.1.1.529 S2P probe showed mostly trimeric particles with shapes similar to other S2P probes.

**S1 Table.** Plasmids from this study and their Addgene accession numbers.

| Plasmid name | Addgene # |
| --- | --- |
| pVRC8400-SARS-CoV-2-S2P-AVI | 160474 |
| pVRC8400-SARS-CoV-2-S2P-D614G-AVI | 176324 |
| pVRC8400-SARS-CoV-2-S2P-B.1.1.7-AVI | 176325 |
| pVRC8400-SARS-CoV-2-S2P-B.1.351-AVI | 176326 |
| pVRC8400-SARS-CoV-2-S2P-P.1-AVI | 176327 |
| pVRC8400-SARS-CoV-2-S2P-B.1.429-AVI | 176328 |
| pVRC8400-SARS-CoV-2-S2P-B.1.526-S477N-AVI | 176329 |
| pVRC8400-SARS-CoV-2-S2P-B.1.526-E484K-AVI | 176330 |
| pVRC8400-SARS-CoV-2-S2P-B.1.617-E154K-L452R-E484Q-D614G-P681R-AVI | 176331 |
| pVRC8400-SARS-CoV-2-S2P-B.1.617-G142D-E154K-L452R-E484Q-D614G-P681R-Q1071H-H1101D-AVI | 176332 |
| pVRC8400-SARS-CoV-2-S2P-B.1.617.1-AVI | 176333 |
| pVRC8400-SARS-CoV-2-S2P-B.1.617.2-AVI | 176334 |
| pVRC8400-SARS-CoV-2-S2P-AY.1-AVI | 176335 |
| pVRC8400-SARS-CoV-2-S2P-B.1.618-AVI | 176336 |
| pVRC8400-SARS-CoV-2-S2P-C.37-AVI | * |
| pVRC8400-SARS-CoV-2-S2P-B.1.621-AVI | * |
| pVRC8400-SARS-CoV-2-S2P-B.1.1.529-AVI | * |
| pVRC8400-SARS-CoV-2-NTD-AVI | 160475 |
| pVRC8400-SARS-CoV-2-NTD-B.1.1.7-AVI | 176424 |
| pVRC8400-SARS-CoV-2-NTD-B.1.351-AVI | 176426 |
| pVRC8400-SARS-CoV-2-NTD-P.1-AVI | 176429 |
| pVRC8400-SARS-CoV-2-NTD-W152C-AVI | 176431 |
| pVRC8400-SARS-CoV-2-NTD-T95I-D253G-AVI | 176432 |
| pVRC8400-SARS-CoV-2-NTD-E154K-AVI | 176433 |
| pVRC8400-SARS-CoV-2-NTD-G142D-E154K-AVI | 176434 |
| pVRC8400-SARS-CoV-2-NTD-B.1.617.1-AVI | 176435 |
| pVRC8400-SARS-CoV-2-NTD-B.1.617.2-AVI | 176436 |
| pVRC8400-SARS-CoV-2-NTD-AY.1-AVI | 176437 |
| pVRC8400-SARS-CoV-2-NTD-B.1.618-AVI | 176438 |
| pVRC8400-SARS-CoV-2-NTD-C.37-AVI | * |
| pVRC8400-SARS-CoV-2-NTD-B.1.621-AVI | * |
| pVRC8400-SARS-CoV-2-NTD-B.1.1.529-AVI | * |
| pVRC8400-SARS-CoV-2-RBD-AVI | 160476 |
| pVRC8400-SARS-CoV-2-RBD-N501Y-AVI | 176439 |
| pVRC8400-SARS-CoV-2-RBD-K417N-E484K-N501Y-AVI | 176440 |
| pVRC8400-SARS-CoV-2-RBD-K417T-E484K-N501Y-AVI | 176441 |
| pVRC8400-SARS-CoV-2-RBD-L452R-AVI | 176442 |
| pVRC8400-SARS-CoV-2-RBD-S477N-AVI | 176443 |
| pVRC8400-SARS-CoV-2-RBD-E484K-AVI | 176444 |
| pVRC8400-SARS-CoV-2-RBD-L452R-E484Q-AVI | 176445 |
| pVRC8400-SARS-CoV-2-RBD-L452R-T478K-AVI | 176446 |
| pVRC8400-SARS-CoV-2-RBD-K417N-L452R-T478K-AVI | 176447 |
| pVRC8400-SARS-CoV-2-RBD-C.37-AVI | * |
| pVRC8400-SARS-CoV-2-RBD-B.1.621-AVI | * |
| pVRC8400-SARS-CoV-2-RBD-B.1.1.529-AVI | * |
| pVRC8400-SARS-CoV-2-RBD-SD1-WA1-AVI | 176448 |
| pVRC8400-SARS-CoV-2-RBD-SD1-N501Y-A570D-AVI | 176449 |
| pVRC8400-SARS-CoV-2-RBD-SD1-K417N-E484K-N501Y+A76-AVI | 176450 |
| pVRC8400-SARS-CoV-2-RBD-SD1-K417T-E484K-N501Y-AVI | 176451 |
| pVRC8400-SARS-CoV-2-RBD-SD1-L452R-AVI | 176452 |
| pVRC8400-SARS-CoV-2-RBD-SD1-S477N-AVI | 176453 |
| pVRC8400-SARS-CoV-2-RBD-SD1-E484K-AVI | 176454 |
| pVRC8400-SARS-CoV-2-RBD-SD1-L452R-E484Q-AVI | 176455 |
| pVRC8400-SARS-CoV-2-RBD-SD1-L452R-T478K-AVI | 176456 |
| pVRC8400-SARS-CoV-2-RBD-SD1-K417N-L452R-T478K-AVI | 176457 |
| pVRC8400-SARS-CoV-2-RBD-SD1-C.37-AVI | * |
| pVRC8400-SARS-CoV-2-RBD-SD1-B.1.621-AVI | * |
| pVRC8400-SARS-CoV-2-RBD-SD1-B.1.1.529-AVI | * |

\* Addgene access codes for these constructs are in the process of being obtained.

